# The effect of gaze on EEG measures of multisensory integration in a cocktail party scenario

**DOI:** 10.1101/2023.08.23.554451

**Authors:** Farhin Ahmed, Aaron R. Nidiffer, Edmund C. Lalor

**Author notes:** **Correspondence: Edmund C. Lalor, E-mail address**.

## Abstract

Seeing the speaker’s face greatly improves our speech comprehension in noisy environments. This is due to the brain’s ability to combine the auditory and the visual information around us, a process known as multisensory integration. Selective attention also strongly influences what we comprehend in scenarios with multiple speakers – an effect known as the cocktail-party phenomenon. However, the interaction between attention and multisensory integration is not fully understood, especially when it comes to natural, continuous speech. In a recent electroencephalography (EEG) study, we explored this issue and showed that multisensory integration is enhanced when an audiovisual speaker is attended compared to when that speaker is unattended. Here, we extend that work to investigate how this interaction varies depending on a person’s gaze behavior, which affects the quality of the visual information they have access to. To do so, we recorded EEG from 31 healthy adults as they performed selective attention tasks in several paradigms involving two concurrently presented audiovisual speakers. We then modeled how the recorded EEG related to the audio speech (envelope) of the presented speakers. Crucially, we compared two classes of model – one that assumed underlying multisensory integration (AV) versus another that assumed two independent unisensory audio and visual processes (A+V). This comparison revealed evidence of strong attentional effects on multisensory integration when participants were looking directly at the face of an audiovisual speaker. This effect was not apparent when the speaker’s face was in the peripheral vision of the participants. Overall, our findings suggest a strong influence of attention on multisensory integration when high fidelity visual (articulatory) speech information is available. More generally, this suggests that the interplay between attention and multisensory integration during natural audiovisual speech is dynamic and is adaptable based on the specific task and environment.

## Introduction

Interpersonal communication heavily relies on successful interpretation of spoken language. While speech is typically perceived through hearing, the ability to observe a speaker’s facial movements and gestures can significantly aid in speech comprehension (Arnold and Hill 2001), particularly in noisy environments (Sumby and Pollack 1954; Ross et al. 2007). This benefit is attributed to a process known as multisensory integration (Stein and Stanford 2008), which is the brain’s ability to integrate information across various senses. In the context of speech, the integration of audio and visual information is thought to occur across two concurrent modes – a correlated mode wherein visual speech provides (temporally correlated) redundant information about the audio speech; and a complementary mode wherein specific visual speech configurations provide unique information about the speaker’s articulatory patterns (Campbell 2008; Peelle and Sommers 2015). Importantly, this latter contribution necessarily depends on having access to detailed visual articulatory information and, as such, likely depends heavily on a listener’s gaze behavior.

Alongside the multisensory integration of audio and visual speech, another phenomenon that greatly affects speech comprehension in noisy environments is selective attention to a particular speaker while blocking out any overlapping competing speakers – famously known as the cocktail-party effect (Cherry 1953). While, most studies investigating this effect considered audio-only scenarios (Ding and Simon 2012b; O’Sullivan et al. 2015; Puvvada and Simon 2017), several have examined the effect in scenarios involving multisensory (i.e., audiovisual) speakers (Zion Golumbic et al. 2013; O’Sullivan et al. 2019). These studies have generally shown that access to visual speech results in a better neural representation of that speech and thus improved success in solving the cocktail party problem (i.e., improved selective attention) leading to better speech comprehension. However, less effort has been dedicated to understanding how selective attention influences the multisensory integration process itself.

The majority of work that has been done on how attention and multisensory integration interact with each other has used non-speech stimuli such as beeps and flashes (Burg et al. 2008; 2011; Talsma, Doty, and Woldorff 2007). And when speech has been used, it has tended to involve fairly simple stimuli such as syllables, isolated words, short segments of speech (Driver 1996; Senkowski et al. 2008; Matusz et al. 2015), including incongruent syllables (Mcgurk and Macdonald 1976) that lead to illusory speech percepts (Alsius et al. 2005; Navarra et al. 2010; Alsius et al. 2014). However, our everyday speech communication is not limited to individual syllables or words. Rather, it involves a continuous and interconnected stream of meaningful words, accompanied by a range of facial and articulatory movements that are temporally correlated with and that complement the ongoing acoustic information (Chandrasekaran et al. 2009; Campbell 2008; Peelle and Sommers 2015). More work is therefore needed to understand the interaction between attention and multisensory integration in the context of natural, continuous audiovisual speech, and particularly, how the correlated and complementary modes of audiovisual integration are affected by attention.

A recent study from our group attempted to explore this question. Specifically, we found that selective attention plays a crucial role in the integration of natural, continuous auditory and visual speech (Ahmed et al. 2023). In that study, we derived an electrophysiological index of multisensory integration using EEG and found evidence for the multisensory enhancement of speech only when that speech was attended. However, that study left several questions unanswered. First, in that experiment, participants were always looking directly at the audiovisual speaker they were attending or ignoring, representing a common real-life scenario where both the correlated and complementary modes of audiovisual speech processing can be utilized. However, in real-life situations, there are often instances where we need to covertly attend to an audiovisual speaker located in our peripheral vision without directly fixating on them, such as when monitoring the environment or avoiding potentially awkward social interactions. In these cases, since we cannot directly look at the face of the audiovisual speaker, the detailed articulatory gestures of that speaker are less available to us and the temporally correlated mode of multisensory integration likely dominates any audiovisual integration. Second, in the previous study, the distractor was always presented in the audio-only modality whereas in real-life, we are frequently faced with distractors in the audiovisual modality, requiring us to suppress competing audio *and* visual information. How attention modulates audiovisual integration across the two scenarios with contrasting gaze behavior and a competing audiovisual distractor is an interesting question that we aim to answer in the present work. Another goal of the present work is that of replication. Specifically, the previous study used a between-subjects design, in the sense that the EEG models employed to index multisensory integration were trained on one group of participants and then tested on a second group. As such, here, we aim to replicate our previous findings using a within-subject design.

To summarize, the goal of the present study is to investigate whether the observed influence of attention on audiovisual speech integration as reported in Ahmed et al. 2023 remains consistent across scenarios where people either do (direct gaze) or do not (peripheral vision) have access to detailed visual articulatory cues. More specifically, we aim to answer the following question: does attention continue to influence multisensory integration, even when the intricate articulatory facial cues are inaccessible, although the temporally synchronized visual cues are still present? To answer this question, we have designed two experiments – one requiring participants to attend to an audiovisual speaker using their peripheral vision (thereby having limited access to the detailed articulatory movements of the speaker while retaining a sense of that speaker’s temporal dynamics; experiment 1), and another that allows participants to look directly at the face of the audiovisual speaker they are attending to (thereby having full access to both temporal dynamics and articulatory details; experiment 2). We recorded high density EEG from the participants as they undertook these experiments and, following previous research (Crosse et al. 2015; 2016a; Ahmed et al. 2023), we indexed multisensory integration by modeling that recorded EEG based on either assuming an underlying multisensory process (AV) or assuming two independent unisensory processes (A+V). In doing so, we found strong evidence for the influence of attention on multisensory integration in conditions where participants had access to the articulatory details (i.e., when they were directly looking at the speaker’s face), but no evidence for such an interaction in the absence of those articulatory details (i.e., when the speaker was viewed in the visual periphery).

## Materials and Methods

### Participants

16 participants (6 males, age range 18 - 29 years, mean 21.31 years) took part in experiment 1 and 16 participants (5 males, age range 18 - 35 years, mean age 22.4 years) took part in experiment 2. One participant in experiment 2 was excluded because of an inadequate amount of data, resulting in a dataset of 15 participants in experiment 2. 7 participants took part in both experiments, with an average gap of 3 months between the two sessions. Each participant provided written informed consent beforehand. All participants were native English speakers, were free of any neurological impairments, had self-reported normal hearing, and had normal or corrected-to-normal vision. Both experiments were approved by the Research Subjects Review Board at the University of Rochester.

### Stimuli and Procedure

The speech stimuli for both experiments were taken from a set of video recordings featuring a well-known male speaker. Each video was cropped into an elliptical shape with the speaker’s head, shoulders and chest centered in the shape with no background movement. Then two such instances of this same speaker (but uttering different content), one on the left and another on the right were combined into a single video file, which served as the audiovisual stimulus for both experiments (Fig. 1). Video editing was done using Adobe After Effects (Adobe Inc.). The linguistic content consisted of current topics related to politics, economics, environment, education etc. and the language was colloquial American English. The audio signals were convolved with a head related transfer function (taken from the MIT KEMAR database, available to download at https://sound.media.mit.edu/resources/KEMAR.html), making them sound like one speech stream was coming from 15° to their left and the other was coming from 15° to their right. The intensity of each soundtrack was normalized by its root mean square using MATLAB (Mathworks). The video and audio files were then merged using the VirtualDub video editor. Because of the way in which we planned to model the EEG data (please see below), we constructed audiovisual stimuli that were either congruent, in which case, the audio was dubbed to its original video or incongruent, in which case, the audio was dubbed to a mismatched video.

**Figure 1:**
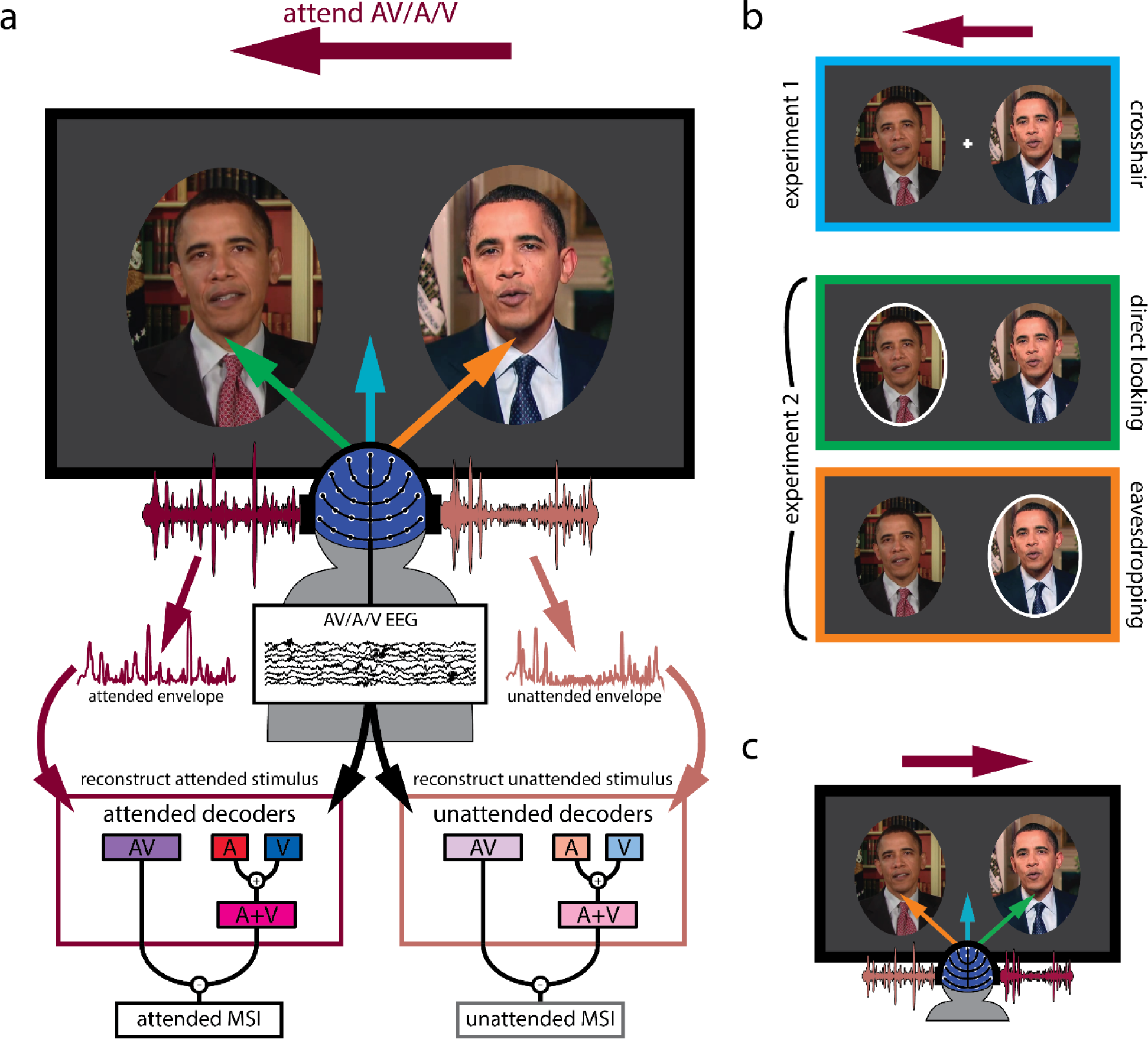
Experimental design. **a.** General outline of both experiments. Participants were presented with two audiovisual speakers, one on the left and another on the right. The deep magenta arrow indicates the speaker (left or right) they had to attend while the blue, green and orange arrows indicate their gaze behavior. They either attended to the audio-only (A), visual-only (V) or audiovisual (AV) modality of the speaker indicated by the text cue. Both attended and unattended AV and (A + V) decoder models were derived from their EEG recordings. As described in the text, these decoders sought to reconstruct an estimate of the speech envelope from the corresponding EEG. Multisensory integration was then quantified as the difference between reconstruction accuracy (Pearson’s r between the actual and the reconstructed envelope of the audiovisual speaker) using the AV decoder and the (A + V) decoder. **b.** Experimental conditions in detail. The deep magenta arrow placed on top of the panels corresponds to attentional cue, indicating that participants were attending to the left stimulus in this <u>example</u>. In experiment 1, participants fixated on a white crosshair placed in the middle of the computer screen (crosshair condition, first panel). In experiment 2, participants either looked at the left speaker that also coincided with the cued attentional side (direct looking condition, middle panel) or at the right speaker marked by the white ellipse while still attending to the left speaker (eavesdropping condition, third panel). Left and right attentional sides were counterbalanced. **c.** Attentional side was counterbalanced. Another example is shown where participants are cued to attend to the right speaker, indicated by the deep magenta arrow. Once again, the blue arrow indicates crosshair fixation, while the green arrow indicates directly looking at the to-be-attended speaker (this time the right speaker) and the orange arrow indicates directly looking at the to-be-ignored speaker (this time the left speaker).

During the experiment, participants viewed these videos as they sat comfortably in a dimly-lit acoustically-shielded booth with their head situated in a chin rest. They watched the videos on a computer monitor in front of them and listened to the corresponding audio track via headphones (Sennheiser HD650). As explained below (in the Modeling EEG responses to natural speech using stimulus reconstruction section), our general analysis strategy involved relating the speech stimuli to the resulting EEG using “decoder” models that are fit to either multisensory AV stimuli (AV decoders) or derived from separate unisensory stimuli (A+V decoders). Moreover, our strategy required us to be able to do this for both attended and unattended stimuli. As such, we needed to present participants with attended and unattended multisensory (AV) stimuli, as well as attended and unattended unisensory (A and V) stimuli. This latter point was the reason why we made the incongruent stimuli mentioned above (technically the stimuli were always multisensory, but having incongruent audio and visual speech meant we could fit unisensory EEG models to either the audio or visual speech, as they were uncorrelated with each other). With those goals in mind, the specific experiments proceeded as follows. Prior to each trial, participants were given an arrow cue to attend to the left or right side (balanced across all conditions) and a text cue for which modality to attend: both auditory and visual (AV) modalities of a congruent AV stimulus, the auditory-alone (A), or the visual-alone (V) modality of an incongruent AV stimulus. Again, it is important to remember that during auditory-alone and visual-alone attention trials, the speech in the other modality was incongruent with the target allowing us to fit a unisensory EEG decoding model by avoiding obligatory multisensory effects driven by temporal correlation (Nidiffer et al. 2018; Maddox et al. 2015).

As mentioned above, there were two experiments involving different participant groups (although 7 participants did both experiments). The main difference between the experiments was the location in which participants were instructed to fixate. Specifically, in experiment 1, participants fixated on a crosshair between the two faces (crosshair fixation; Fig. 1a blue arrow; 1b top panel). Meanwhile, no fixation crosshair was present in experiment 2. Instead, participants were instructed to directly look at a speaker’s face, cued by a white ellipse placed around the speaker (Fig. 1b, middle and bottom panels). However, importantly, in this experiment, the targets of attention and gaze could be different; participants either looked directly at the to-be-attended speaker (direct looking; Fig. 1a green arrow; 1b middle panel) or looked toward the to-be-ignored speaker, while actually attending to the other speaker (eavesdropping; Fig. 1a orange arrow; 1b bottom panel). In experiment 1, there were 20 trials per condition (3 attending modalities: audiovisual, audio-only, visual-only), resulting in a total of 60 trials. Experiment 2 had 14 trials per condition (3 attending modalities as before × 2 gaze directions: direct looking vs. eavesdropping), resulting in 84 trials in total. Each trial was 1-min long. Stimulus presentation was controlled using Presentation software (Neurobehavioral Systems).

To ensure that participants were engaging with the appropriate visual content, some visual targets were incorporated into the videos. In the case where they were attending to a congruent audiovisual speaker, the targets took the form of brief interruptions or glitches consisting of temporary (tens of milliseconds) warping of the speaker’s face, freezing video frames or repeated movements of the mouth or head. In the case where they were attending to the visual speech modality alone (i.e., when presented with an incongruent video), the targets consisted of some gradual slow-motion (frame rate of the video reduced to 70% of original rate) events lasting for about 2 sec. The reason for the two different types of targets was as follows. We reasoned that, during AV attention, participants would primarily concentrate on the content of the audio speech. And we wanted to motivate participants to care about inconsistencies between that audio speech and the accompanying visual speech. So, we inserted the short temporal glitches (described above) that they were required to respond to. Meanwhile, in the V-only condition, we worried that attending for glitches might mean that participants were not attempting to read the visual stimulus as speech per se, but rather watching out for general visual discontinuities. As such, in this case, the targets consisted of slow-motion events. Our rationale was that this would encourage participants to carefully monitor the mouth and face dynamics (i.e., to attempt to speech read the videos) to maximize their chances of detecting these subtle slowdowns. Indeed, careful behavioral pilot testing was carried out before the experiments to determine the parameters of the glitches and the slow-motion events so that they were challenging enough to be detected when participants were actively engaged with the visual content, but not so easy to detect that they would simply pop out. Both kinds of visual targets appeared randomly anywhere between 1 and 6 times per one-minute trial. Participants had to press the spacebar as soon as they detected a target. A response within 2 secs of a target glitch event or within 4 secs of a slow-motion event onset was recorded as a hit, while responses outside this boundary were recorded as false alarms.

To make sure participants were engaging with the appropriate audio in the cases where they were attending to a congruent audiovisual speaker or the audio speech modality alone of an incongruent audiovisual speaker, they answered two comprehension questions, each with four possible answers about the attended auditory speech stream after each trial. The tasks were consistent across both experiments. Please note that participants were not asked to detect visual targets in the attend A-only trials (since we did not want them to engage with the visual content) and no comprehension questions were asked after the attend V-only trials (since we did not want them to engage with the auditory content). Table 1 presents a comprehensive overview of all the conditions.

**Table 1:** Experimental conditions. Subscripts are chosen according to gaze behavior; c for central fixation, d for direct looking at the attended speaker, e for eavesdropping/looking away from the attended speaker. Q/A stands for comprehension question answering.

| Condition | Fixation | Attended modality | AV correspondence | Task |  |
| --- | --- | --- | --- | --- | --- |
| AV<sub>c</sub> | Crosshair | AV | Congruent | Detect glitches, | Experiment 1<br>(N = 16) |
| A<sub>c</sub> | Crosshair | A | Incongruent | Answer Q/A |  |
| V<sub>c</sub> | Crosshair | V | Incongruent | Detect slow motion |  |
| AV<sub>d</sub> | Attended speaker | AV | Congruent | Detect glitches, Q/A | Experiment 2<br>(N = 15) |
| A<sub>d</sub> | Attended speaker | A | Incongruent | Answer Q/A |  |
| V<sub>d</sub> | Attended speaker | V | Incongruent | Detect slow motion |  |
| AV<sub>e</sub> | Ignored speaker | AV | Congruent | Detect glitches, Q/A |  |
| A<sub>e</sub> | Ignored speaker | A | Incongruent | Answer Q/A |  |
| V<sub>e</sub> | Ignored speaker | V | Incongruent | Detect slow-motion |  |

### EEG recording and preprocessing

In both experiments, EEG data were recorded at a sampling rate of 512 Hz from 128 scalp electrodes and two mastoid electrodes using a BioSemi ActiveTwo system. Subsequent preprocessing was conducted offline in MATLAB. The data were bandpass filtered between 0.3 and 30 Hz, down sampled to 64 Hz and referenced to the average of the mastoid channels. To detect channels with excessive noise, we used EEGLAB to determine which channels showed EEG with unusual values in terms of its spectrogram, probability distribution and kurtosis (Delorme and Makeig 2004). Channels were also marked as noisy if their variance exceeded three times the average variance of all channels. Noisy channels were recalculated using spline interpolation in EEGLAB. Afterwards, the EEG data were z-scored.

### Eye tracking

To ensure that participants were looking at the appropriate content, eye movements were monitored throughout the experiments. Eye data was recorded using Tobii Pro X3-120 eye tracker. The eye tracker was mounted at the base of the computer monitor and data were collected at a sampling rate of 120 Hz. Calibration was carried out every 15 trials using nine target points distributed over the entire screen.

### Envelope extraction

Given that most people rely primarily on acoustic cues to understand speech, our analysis strategy focused on relating EEG to the speech acoustics and assessing how that relationship varied with multisensory input and with attention. Specifically, we aimed to relate the EEG to the acoustic envelopes of the speech stimuli – an approach that has been widely used in previous research (Aiken and Picton 2008; Lalor and Foxe 2010; Ding and Simon 2014; Ahissar et al. 2001; Peelle, Gross, and Davis 2013). To obtain the acoustic envelopes of the speech stimuli from each trial, we first bandpass filtered those stimuli through a gammatone filter bank into 128 logarithmically spaced frequency bands between 100 Hz and 6500 Hz. Next, we used the Hilbert transform to obtain the envelope for each of the 128 frequency bands, and finally, we determined the broadband envelope by averaging the narrowband envelopes across all 128 bands. Later, the envelopes were z-scored.

In the case of the incongruent audiovisual stimuli, when participants were attending to the auditory modality alone (i.e., they couldn’t see the face matching the audio speech), the envelope of the audio speech was extracted. When participants were attending to the visual modality alone (i.e., they couldn’t hear the audio speech matching the visual speech), the acoustic envelope of the unheard speech corresponding to the visual speech that was presented on the monitor was extracted. Following previous work (Crosse et al. 2015; 2016a; Ahmed et al. 2023), and as discussed in more detail below, this was done to account for visual cortical responses over occipital scalp to motion in the video that correlates with the unheard audio envelope.

### Modeling EEG responses to natural speech using stimulus reconstruction

As mentioned, our primary dependent measure was how well the acoustic envelope of the speech stimuli was reflected in the EEG – with the goal of assessing how that measure changed with attention and multisensory input. To derive this measure, we used linear decoding models (temporal response functions) to reconstruct an estimate of the audio speech envelope from the corresponding multichannel EEG data (as described in Crosse et al. 2016b). Specifically, we fit the following model:

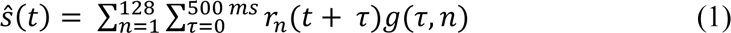

where *ŝ*(*t*) is the reconstructed envelope, *r*_*n*_(*t* + *τ*) is the EEG response at channel n, *τ* is the relative time lag between the ongoing stimulus and the EEG data, and *g*(*τ*, *n*) is the linear decoder for the corresponding channel and time lag. The time lags *τ* ranged from 0 to 500 ms post-stimulus. The decoder *g*(*τ*, *n*) was obtained using ridge regression as follows:

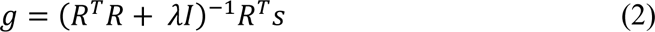

where *R* is the lagged time series of the EEG data, *λ* is a regularization parameter (known as the ridge parameter), *λ* is the identity matrix and *ŝ* is the speech envelope. To fit the decoders, we employed a leave-one-trial-out cross-validation approach to determine the optimal λ value within a range of 10^−6^, 10^−4^, …, 10^30^. This λ value was chosen to maximize the correlation between the estimated reconstructed speech envelope, ŝ(t) and the actual speech envelope s(t), while avoiding overfitting to the training data. Using this approach, we optimized modality-specific decoders using the EEG and the acoustic envelopes from trials where participants attended the A, V, or AV modalities of the presented speech streams. The (A+V) decoder that represents a linear of combination of the two independent unisensory processes was trained on the summed covariance matrices of the unisensory responses (EEG data corresponding to attend A-only and attend V-only conditions respectively). Importantly, the (A+V) decoder was validated by confirming that it could reconstruct the envelope of the speech presented during the AV condition, using EEG from that condition. This was done because we were interested to see whether the data obtained during a multisensory (AV) task is better modeled as an AV process compared to the independent summation of two unisensory processes. Again, the λ parameter for the (A+V) decoder was optimized using a leave-one-trial-out cross-validation approach that maximized the correlation between the reconstructed and the actual speech envelope of the congruent AV stimuli. The λ optimization was done separately for each participant and for each condition. We used mTRF-toolbox-2.3 (available to download at https://github.com/mickcrosse/mTRF-Toolbox/releases/tag/v2.3) to fit the decoding models.

As there were 2 simultaneous congruent/incongruent audiovisual speech streams in each trial (one attended, another unattended), we trained both attended and unattended AV and (A+V) decoders for each participant, where linear regression was performed between the EEG data and the attended or unattended speech respectively. In total, we trained 24 decoders: 4 modalities (AV, A, V, A+V) × 2 attentional states (attended, unattended) × 3 gaze conditions (crosshair, c; direct looking, d; eavesdropping, e). Table 2 presents a comprehensive overview of the attended/unattended decoders fitted from each condition.

**Table 2:** List of decoders used in our analyses, along with the condition each decoder was fit from, and the speech envelope reconstructed using each decoder. As in Table 1, subscripts are chosen according to gaze behavior; c for central fixation, d for direct looking at the attended speaker, e for eavesdropping/looking away from the attended speaker. The AV and (A+V) decoders are bolded to emphasize the point that these are the primary decoders used in this study to quantify multisensory integration as well as for testing our main hypothesis, while the other decoders are fit in order to derive the (A+V) decoders.

| Decoders | Fixation | Decoders fit from Condition | Envelope reconstructed (using the decoder) |  |
| --- | --- | --- | --- | --- |
| **AV<sub>c</sub>** | **Crosshair** | **AV<sub>c</sub>** | **AV<sub>c</sub>** | Experiment 1<br>(N = 16) |
| **(A+V)<sub>c</sub>** | **Crosshair** | **A<sub>c</sub>, V<sub>c</sub>** | **AV<sub>c</sub>** |  |
| A<sub>c</sub> | Crosshair | A<sub>c</sub> | A<sub>c</sub>, AV<sub>c</sub> |  |
| V<sub>c</sub> | Crosshair | V<sub>c</sub> | V<sub>c</sub>, AV<sub>c</sub> |  |
| **AV<sub>d</sub>** | **Attended speaker** | **AV<sub>d</sub>** | **AV<sub>d</sub>** | Experiment 2<br>(N = 15) |
| **(A+V)<sub>d</sub>** | **Attended speaker** | **A<sub>d</sub>, V<sub>d</sub>** | **AV<sub>d</sub>** |  |
| A<sub>d</sub> | Attended speaker | A<sub>d</sub> | A<sub>d</sub>, AV<sub>d</sub> |  |
| V<sub>d</sub> | Attended speaker | V<sub>d</sub> | V<sub>d</sub>, AV<sub>d</sub> |  |
| **AV<sub>e</sub>** | **Ignored speaker** | **AV<sub>e</sub>** | **AV<sub>e</sub>** |  |
| **(A+V)<sub>e</sub>** | **Ignored speaker** | **A<sub>e</sub>, V<sub>e</sub>** | **AV<sub>e</sub>** |  |
| A<sub>e</sub> | Ignored speaker | A<sub>e</sub> | A<sub>e</sub>, AV<sub>e</sub> |  |
| V<sub>e</sub> | Ignored speaker | V<sub>e</sub> | V<sub>e</sub>, AV<sub>e</sub> |  |

### Quantifying multisensory integration

Our aim was to explore how the impact of attention on multisensory integration varies depending on the amount of visual speech detail available to the participants. We achieved this by measuring multisensory integration (MSI) using the additive model criterion (Stein and Meredith 1993). We then examined how this measure varied across attentional states in settings where participants had limited access to the facial characteristics of the target speaker (experiment 1) vs where they had full access (experiment 2). The additive model criterion defines multisensory integration as the difference between cortical responses to multisensory stimuli (AV) and the sum of the responses to unisensory stimuli (A + V). To that end, as discussed in the previous section, we trained separate linear models/decoders for attended as well as unattended audio (A), visual (V), and audiovisual (AV) speech modalities using corresponding EEG data. The algebraic sum of the A and V decoders was then computed to obtain an (A+V) decoder which represents the independent processing of each sensory modality.

The audio envelope of the congruent audiovisual speech was reconstructed using both AV and (A+V) decoders in both experiments according to equation (1). Multisensory integration was then quantified as:

Experiment 1 (crosshair fixation)

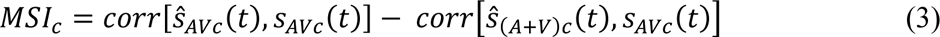

Experiment 2 (direct looking at the target)

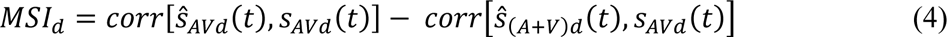

Experiment 2 (looking at the ignored speaker while eavesdropping on the target)

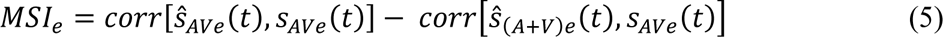

And, again, these MSI values were calculated for stimuli that were attended (using all “attended” decoders) and for stimuli that were unattended (using all “unattended” decoders), so that we could quantify the influence of attention on multisensory integration.

### Single-lag analysis

In our main analyses – described above – we fit decoders *g*(*τ*, *n*) to incorporate EEG data across a wide range of time lags from 0 to 500 ms. This choice was made with the goal of incorporating all EEG changes related to a unit change in the envelope (i.e., one might expect that a change in the acoustic envelope at any point in time would be seen in the EEG between 0 and 500 ms later). However, it would also be interesting to explore the possibility of differences in how attention and multisensory integration interact across time lags. For example, previous research has suggested that multisensory integration can happen automatically and independent of attention (i.e., pre-attentively) based on shared temporal properties (Van der Burg et al. 2011; Atilgan et al. 2018). Whereas we might expect a greater influence of attention on multisensory integration at longer latencies given previous research on cocktail party attention (Power et al. 2012; Patel et al. 2022)

To examine this possibility, and more generally, to examine how the interplay between attention and multisensory integration unfolds over time, we implemented a time-resolved analysis by training an AV and (A+V) decoder in each condition at every single time lag within the range of −500 to 500 ms (instead of integrating across these time lags). With a sampling frequency of 64 Hz, this approach produced a total of 65 separate time lags at an interval of 15.625 ms and consequently, 65 distinct AV decoders and 65 separate (A+V) decoders in each condition. These decoders were then used to reconstruct the relevant audiovisual speakers in both experiments, giving us a sense of how attention and multisensory integration interact at each time lag within our range.

### Statistical analysis

To test that the decoding models could reconstruct the relevant speech envelopes significantly better than chance, nonparametric permutation tests were performed (Combrisson and Jerbi 2015). For each participant, a null distribution, consisting of 1000 Pearson’s *r* values, was generated by shuffling the stimulus-EEG pair between trials and calculating a null model with each shuffle. Then the average of those 1000 values was obtained for each participant. Group-level Wilcoxon signed-rank testing was then performed between these null correlation coefficients and the actual trial-averaged correlation coefficients. This procedure was repeated for each decoder and for each attentional state in both experiment 1 and 2. The threshold for above chance performance was *p* = 0.05 for each test. Within the same groups of participants, paired Wilcoxon signed-rank testing was used for statistical comparison across conditions (attended vs unattended) and across decoders (AV vs A+V). However, since different groups of participants took part in experiment 1 and 2, whenever a direct comparison was made between the two experiments, unpaired Wilcoxon rank-sum testing was performed (although seven participants were common to both experiments, for simplicity, we treated them as unpaired at the cost of losing some statistical power, but being more conservative with our statistical test). False discovery rate (FDR) method (Benjamini and Hochberg 1995) was used for correcting the *p*-values in the single-lag analysis where multiple comparisons were made. Numerical values are reported as mean ± standard deviation.

### Linear mixed effects modeling

To supplement our within-experiment analyses, we wanted to investigate effects across the combined results of the two experiments while accounting for participant variability and the shared variance of the participants who volunteered in both experiments (n = 7). To do so, we fit a series of linear mixed-effects (LME) models to explain different response variables from some variable of interest and other potentially explanatory control variables (e.g., experiment effects, subject random effects). Generally, for each comparison, we fit two models: one including the variable of interest as well as any control variables (full model) and another including just the control variables (reduced or null model). For example, to examine the effect of gaze (gz) on comprehension score (qa), we fit:

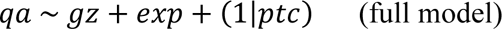

and

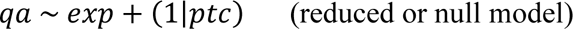

where the variable of interest is gaze (gz), while controlling for the differences between experiments (exp) and overall participant variability (ptc). In all our LME results, we reported the regression coefficient (slope of the regression line explaining the relationship between predictor and response variables, β) and the *p* value. The fitted models were also compared using the Log Likelihood Ratio (LLR) Test that compares the fit of the two models on the data. A positive likelihood ratio provides support for the full model over the null model.

We also use LME models to measure gaze’s effect on speech tracking, multisensory interactions, and attentional benefits. Effects related to speech tracking were taken from the reconstruction accuracies on the AV attended trials. The multisensory benefit was again modeled with our standard AV-(A+V) approach using the data on AV attended trials. Finally, the attentional benefit was modeled by subtracting the attended and unattended reconstruction accuracies during the AV trials. With the help of our single-lag decoder analysis, reconstruction accuracy and thereby our linear mixed-effects models could be resolved over time-lag.

## Results

### Participants deployed attention to the target speech stimuli

As discussed above, the various conditions in our experiments were deliberately designed to enable us to derive attended and unattended AV, A, and V decoders. Again, ultimately, this was to allow us to test how well an attended/unattended AV decoder would perform relative to an attended/unattended (A+V) decoder in reconstructing the speech envelope of an attended/unattended (congruent) audiovisual speech stimulus. As such, we incorporated both visual and auditory tasks into our different conditions to motivate participants to attend to a particular stimulus while ignoring the others (see Table 1).

The d-prime scores suggest that participants were able to detect visual targets in both experiments (Fig. 2a; d-prime scores in different conditions — AV_d_: 3.50 ± 0.82, AV_c_: 2.82 ± 0.64, AV_e_: 2.15 ± 0.61, V_d_: 2.46 ± 0.81, V_c_: 1.54 ± 0.89, V_e_: 1.1 ± 0.55). This alone does not definitively prove that the participants were actively attending to only one visual stimulus (after all, participants were not asked to respond to visual targets in the unattended stream). However, as mentioned earlier, behavioral pilot testing was conducted before implementing the EEG experiments so as to make the targets difficult enough to detect. As such, we contend that these strong d-prime scores provide evidence that participants were attending to the target visual stimuli – and ignoring the non-target visual stimuli – enabling us to fit our attended and unattended decoders.

**Figure 2:**
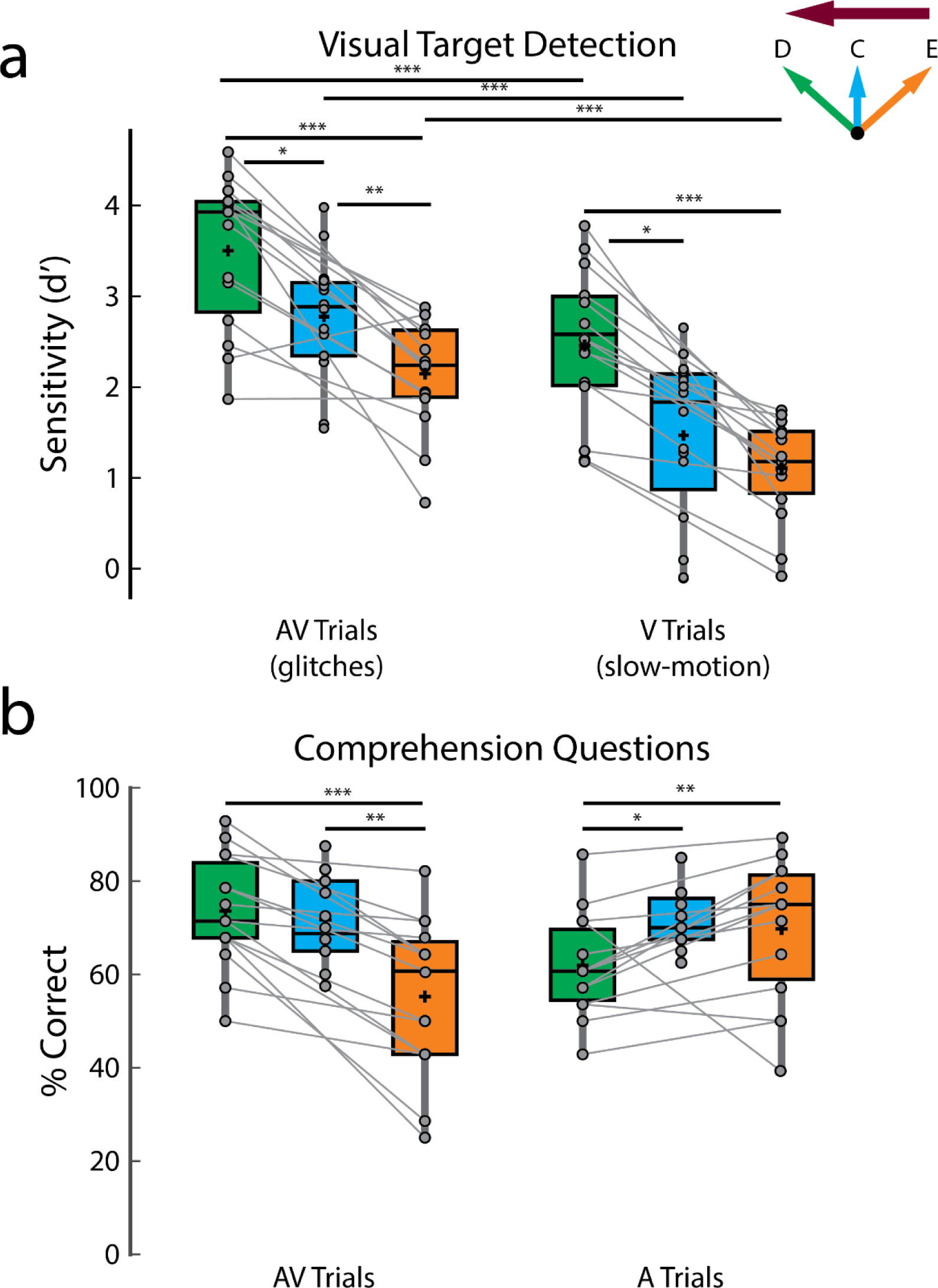
Behavioral performance. **a.** Sensitivity (d’) scores on detecting visual targets. **b.** Scores on answering comprehension questions. The green, blue, and orange colors indicate direct looking (D), crosshair (C) and eavesdropping (E) conditions respectively and the deep magenta arrow indicates the attentional side (can be either left or right). Connected lines are established between green and orange panels, since they belong to the same experiment involving same set of participants (experiment 2) while the blue panel belongs to a separate experiment (experiment 1), involving a different set of participants (except for 7 participants who participated in both experiments). *** *p* < 0.001, ** *p* < 0.01, * *p* < 0.05.

In terms of the effect of gaze behavior on target detection, participants were better at detecting visual targets in the direct looking compared to the eavesdropping conditions (Fig. 2a; d-prime AV_d_ vs AV_e_, *p* = 3.05 × 10^−4^, V_d_ vs V_e_: *p* = 6.10 × 10^−5^; Wilcoxon signed-rank tests) and performance in crosshair fixation generally fell in between (Fig. 2a; d-prime AV_c_ vs AV_d_, *p* = 0.02, AV_c_ vs AV_e_, *p* = 0.0068, V_c_ vs V_d_, *p* = 0.01, although close to significance, V_c_ vs V_e_, *p* = 0.06; unpaired Wilcoxon rank-sum tests, since different participants took part in experiment 1 except for seven participants). Additionally, as described in the methods section, we fit linear mixed-effect (LME) models including all data to investigate the effect of gaze on visual target (glitch or slow-motion) detection scores while controlling for individual and experimental differences. We found that gaze is a strong predictor of glitch detection (β = −0.47; *p* = 4.03 × 10^−9^) and slow-motion detection (β = −0.69; *p* = 1.84 × 10^−29^) with gaze providing an overall better fit for both models compared to the null model without gaze (predicting glitch: LLR = 17.80, *p* = 2.5 × 10^−5^; predicting slow-motion: LLR = 126.80, *p* = 0). These results reflect the general difficulty as gaze moves away from the target of attention – with the direct looking condition being easiest, the crosshair condition being more difficult, and the eavesdropping condition being the most difficult. One additional comment here: we did not attempt to perfectly match the two types of visual target in terms of the difficulty of detecting them. However, for completeness, we note that participants struggled more in detecting the slow-motion targets than the glitches. Interpreting this is not completely straightforward, but one speculation is that it may reflect more difficulty with lipreading in general when the corresponding acoustic speech is unavailable (Fig. 2a; d-prime AV_d_ vs V_d_, *p* = 3.05 × 10^−4^, AV_c_ vs V_c_, *p* = 4.38 × 10^−4^, AV_e_ vs V_e_, *p* = 3.05 × 10^−4^; Wilcoxon signed-rank tests).

In terms of the audio task, participants performed significantly better than the 25% chance level in answering the multiple-choice comprehension questions in all conditions (Fig. 2b; comprehension scores in different conditions — AV_d_ : 73.57 ± 11.91%, *p* = 6.10 × 10^−5^, AV_c_: 71.72 ± 8.88 %, *p* = 6.10 × 10^−5^, AV_e_ : 55.24 ± 16.53 %, *p* = 6.10 × 10^−5^, A_d_ : 62 ± 10.77%, *p* = 6.10 × 10^−5^, A_c_: 72.03 ± 6.53%, *p* = 4.21 × 10^−4^, A_e_ : 69.76 ± 14.65 %, *p* = 6.10 × 10^−5^, Wilcoxon signed-rank tests against 25% chance level). This provides strong evidence of attention to the target speech (and inattention to the non-target speech) based on a long history of research on the cocktail party phenomenon (Cherry 1953). Again, as we mentioned before, this allows us to fit the relevant attended and unattended decoders. Across gaze conditions, the scores on comprehension questions on the AV attending trials largely followed the same pattern as for the visual target detection (Fig. 2b, left panel). Specifically, participants performed better in the direct looking condition compared to eavesdropping (Fig. 2b left panel; AV_d_ vs AV_e_, *p* = 6.10 × 10^−5^, Wilcoxon signed-rank test) and better in the crosshair condition compared to eavesdropping (Fig. 2b left panel; AV_c_ vs AV_e_, *p* = 0.0041, unpaired Wilcoxon rank-sum test). However, there was no significant difference between crosshair and direct looking condition in comprehension scores on AV trials. Interestingly, this general pattern was reversed in comprehension scores on the A attending trials where participants performed worst in the A_d_ condition compared to both the A_c_ and A_e_ conditions (Fig. 2b right panel; comprehension score A_c_ vs A_d_: *p* = 0.0043, A_e_ vs A_d_: *p* = 0.01). This is likely because participants were attending to the auditory speech (played via headphones) of an incongruent AV speaker in the A_c_, A_d_ and A_e_ conditions; however, in the A_c_ and A_e_ conditions, deploying attention away from their gaze site may have helped suppress the uninformative visual stimulus more compared to the A_d_ condition where the incongruent (and as such, uninformative) visual stimulus and the to-be-attended auditory speech are both competing for attention at the same spatial location.

We also used an eye tracker to monitor if participants were fixating on the central crosshair in experiment 1, directly looking at and eavesdropping appropriately on the attended speaker in experiment 2. We found that participants fixated on the instructed speech content most of the time during the 60 sec long trials rarely deviating from their instructed fixation area. Across participants, the percentage of time spent *not* looking at the crosshair in experiment 1 is 2.79 ± 4.07% (averaged across all conditions and trials). In experiment 2, the percentage of time spent *not* looking at the appropriate audiovisual speaker is 5.86 ± 6.4% in the direct looking conditions and 14.49 ± 24.04% in the eavesdropping conditions. Compared to the direct looking conditions, participants tended to deviate more in the eavesdropping condition where they had to look at the ignored audiovisual speaker (*p* = 0.04, Wilcoxon signed-rank test). This is not surprising, given that people can be naturally tempted to look at the speaker they are intending to pay attention to.

### Multisensory integration is strongly modulated by attention when detailed articulatory features are accessed by direct gaze

To investigate multisensory interactions in our experiments, we modeled cortical responses to multisensory (AV) as well as unisensory stimuli (A, V). However, comparing AV tracking to A or V tracking alone is not sufficient to quantify multisensory enhancement since both auditory and visual processes occur during audiovisual speech and their additive combination can benefit speech tracking in the absence of any interactions. Following previous research (Crosse et al. 2015; 2016a; Ahmed et al. 2023), we accounted for this by first summing the unisensory A and V decoders to create (A+V) decoders and then comparing it to the AV model in terms of how well it could reconstruct the speech envelope of the congruent audiovisual speaker. If audiovisual speech processing simply involves the auditory system processing the audio speech and the visual system processing the visual speech, then an (A+V) decoder and an AV decoder should perform equivalently. If they don’t, any difference can be attributed to multisensory integration. We verified that each of these models could predict EEG activity significantly better than chance and that the AV and A+V decoders account for both auditory and visual speech processing (see supplemental figure 1 and text).

For both AV and (A+V) models, the performance of the attended decoders was significantly better than that of the unattended decoders in all gaze conditions (Fig 3a; attended vs unattended reconstruction accuracies — AV_d_: *p* = 1.22 × 10^−4^; (A+V)_d_: *p* = 1.83 × 10^−4^; AV_c_: *p* = 9.35 × 10^−4^; (A+V)_c_: *p* = 5.31 × 10^−4^; AV_e_: *p* = 4.27 × 10^−4^; (A+V)_e_: *p* = 0.0034; Wilcoxon signed-rank tests). This finding is in line with previous studies showing preferential representation of attended speech in the brain over ignored speech (O’Sullivan et al. 2015; Ding and Simon 2012a).

**Figure 3:**
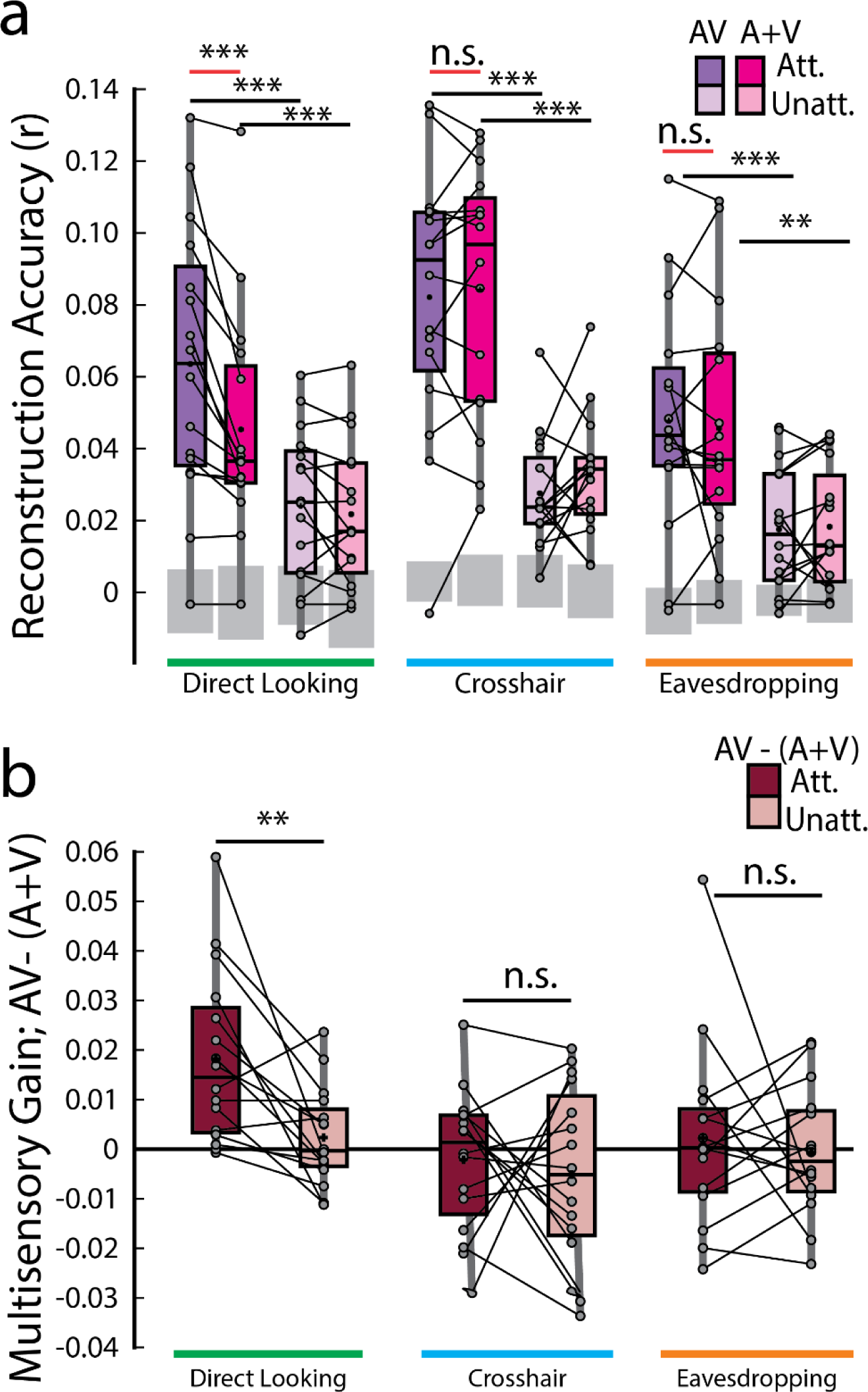
Speech envelope reconstructions. **a.** Envelope reconstruction accuracies averaged across trials for each decoder and gaze condition. Gray boxes indicate chance levels established via permutation testing for each gaze condition. **b.** Multisensory gain (i.e., AV –(A+V) reconstruction accuracies) averaged across trials. Each dot represents an individual participant. *** *p* < 0.001, ** *p* < 0.01, * *p* < 0.05, n.s. stands for not significant.

According to the above-mentioned additive-model criterion, multisensory integration can be measured from the difference between cortical responses to multisensory stimuli and the summed unisensory responses [i.e., AV – (A+V)]. Therefore, the crucial comparison was between the envelope reconstruction accuracies of the AV and (A+V) decoders across gaze conditions. Interestingly, this measure of multisensory integration turned out to be significant *only* in the direct looking condition where participants were directly looking at the audiovisual speaker that they were also paying attention to and in neither of the crosshair or eavesdropping condition (red lines in Fig. 3a; AV_d_ attended vs (A+V)_d_ attended, *p* = 1.22 × 10^−4^; Wilcoxon signed-rank test), suggesting that response to an attended audiovisual speech is not simply the sum of independent audio and visual speech, rather, it includes nonlinear multisensory contributions that is built into the AV model, but not into the (A+V) model. Importantly though, this is evident *only* when the audiovisual speaker is in the direct gaze focus of the participants enabling them to make use of both correlated and complementary visual information. We further investigated whether the multisensory gain index (AV-(A+V)) varies across attended vs unattended speech. Once again, the only time this index varied was in the direct looking condition in experiment 2 (MSI gain attended vs MSI gain unattended, *p* = 0.0015, Wilcoxon signed-rank test; Fig. 3b). This suggests that attention strongly modulates multisensory integration when participants can directly look at their targeted audiovisual speaker.

To more fully explore the impact of gaze on speech tracking and multisensory integration we took a two-pronged approach involving a direct comparison between conditions in experiment 2 and combining data from both experiments in a linear mixed-effects model (see methods). First, we found that the tracking of the attended audiovisual speech was marginally improved in the direct looking condition over the eavesdropping condition (AV_d_ vs AV_e_, *p* = 0.055). Second, we combined data across experiments 1 and 2 into a linear mixed-effects model and predicted attended audiovisual speech tracking from gaze while controlling for experimental and individual differences. We found that gaze is significantly predictive of speech tracking (β = −0.0083; *p* = 0.0023) in the full model and contributes to a better overall fit compared to a null model (LLR = 9.33, *p* = 0.0023). We also found that, in experiment 2, as expected, multisensory integration of the attended audiovisual speech was more pronounced in the direct-looking condition compared to the eavesdropping condition (*p* = 0.0125; Wilcoxon signed-rank test) as well as the crosshair condition (*p* = 0.0015; again, considering unpaired Wilcoxon-rank sum test as we did for behavioral analysis, since different participants took part in those two conditions, except for 7 participants). And again, after combining data across experiments through LME modeling, we found that gaze strongly predicts multisensory integration (β = −0.0085; *p* = 0.0030) over a reduced null model (LLR = 8.81; *p* = 0.0030).

### Exploring the temporal dynamics of multisensory integration

We performed a temporally-resolved analysis to investigate how any multisensory integration effects might vary across time lags. Originally, we had planned to also assess how these multisensory integration effects might interact with attention – again given previous research distinguishing pre-attentive multisensory binding from later integration effects that can be strongly modulated by attention (Van der Burg et al. 2011; Atilgan et al. 2018). However, given that we only found multisensory integration effects for the attended conditions (involving direct looking) and not for any unattended condition, we focused this temporal analysis solely on the attended conditions.

The time-courses of the AV and (A+V) reconstruction accuracies share similar peak pattern across all gaze conditions – one at around 78 ms and another peak at around 250-296 ms. However, the time-courses of the AV and (A+V) decoders closely follow each other in both crosshair and eavesdropping condition while, in line with our earlier result (Fig. 3b), they diverge only in the direct looking condition (Fig. 4b). We found that in the direct looking condition, the AV decoder significantly outperformed the (A+V) decoder across a broad range of time lags between 200 to 500 ms (gray vertical bar in the left panel of Fig. 4b; AV_d_ vs (A+V)_d_, all *p* < 0.05, Wilcoxon-signed rank test, FDR-corrected). This fits with previous studies reporting that cocktail party attention effects tend to manifest at temporal latencies beyond ∼100 ms (Power et al. 2012). It is therefore possible that because of this late attentional modulation, a stronger multisensory integration effect (i.e., AV > (A+V)) becomes apparent only at these longer latencies. As expected, based on our results in the previous section (Fig. 3b), there was no evidence of any multisensory integration effect at any of the time lags for the attended audiovisual speaker in either the crosshair or the eavesdropping condition (middle and right panels of Fig. 4b).

**Figure 4:**
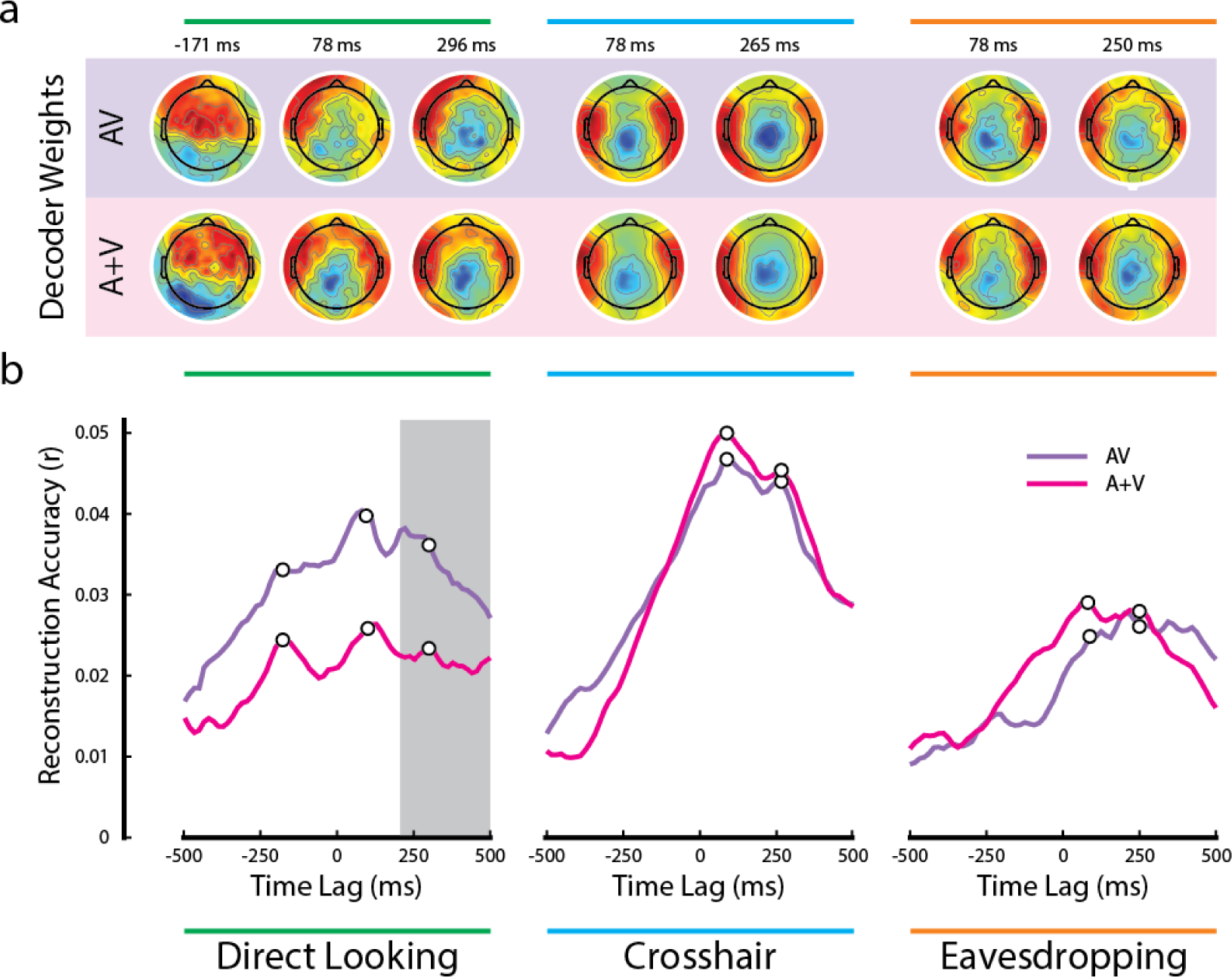
Temporal analysis of multisensory integration. **a.** Decoder weights at the time points corresponding to the white dots in panel b. **b.** Time courses of the AV and (A+V) reconstruction accuracies. The gray bar indicates the time lags when the multisensory effect was significant in the direct looking condition (all p < 0.05; FDR corrected for multiple comparisons).

Notably, there was another distinct peak at a pre-stimulus lag of ∼ −171 ms in both AV and (A+V) reconstruction accuracies that was present in the direct looking condition, but not in the other two conditions. We speculate that this is happening because visual speech signals typically precede the actual production of speech sounds (Schroeder et al. 2008; Van Wassenhove et al. 2005). In the case of natural audiovisual speech, which is what we have in the current study, the correlated visual movements can occur several hundred milliseconds before the sound begins (Chandrasekaran et al. 2009). While the AV-(A+V) effect in the direct looking condition did not reach statistical significance during the early time lags, it is noteworthy that the pre-stimulus peak at −171 ms diverging between AV and (A+V) was observed exclusively in the direct looking condition (Fig. 4b, left panel) and not in the other two gaze conditions. This suggests the possibility of some early multisensory binding occurring at shorter latencies, which is subsequently reinforced by attention at later latencies, ultimately resulting in a multisensory integration effect in our EEG measures. However, we must remain speculative on this point given the lack of any statistical significance at these early latencies. Future work could test this further by endeavoring to improve the SNR of the measures at these latencies by focusing only on this condition and by increasing participant numbers.

Figure 4a shows the channel weights for each decoder at several peak points (marked by the white dots on Fig. 4b). Although the decoder weights may not directly represent the neural generators underlying the observed effects, it is remarkable that the decoder weights maintain similar topographic pattern, particularly in the crosshair and in the eavesdropping conditions. Besides, they all show dominant activities over temporal scalp, suggesting their contribution towards modeling auditory cortical activities.

We again leveraged linear mixed-effects modeling and the single-lag stimulus reconstruction approach to investigate the dynamic influence of gaze on speech tracking, multisensory integration, and attention. For each single time-lag, we fitted a linear model to predict each of these measures from gaze (Figure 5). We found that they can be predicted by gaze over a broad range of time-lags – including many negative lags. Speech tracking during audiovisual trials can be predicted over the broadest range (Fig. 5a): as early as −438 ms pre-stimulus through 156 ms after the stimulus onset and between 203-312 ms post-stimulus. These lags include prominent peaks around the latency visual information becomes available (∼-120ms;Karthik et al. 2022; Schroeder et al. 2008), during early auditory processing (∼100ms) and a late epoch (∼250 ms) that likely reflects a previously reported attentional effect (Power et al. 2012). Enigmatically, we found this relationship extends well before the onset of the speech, far beyond what we would conservatively estimate to be the effects of smearing (see below).

**Figure 5:**
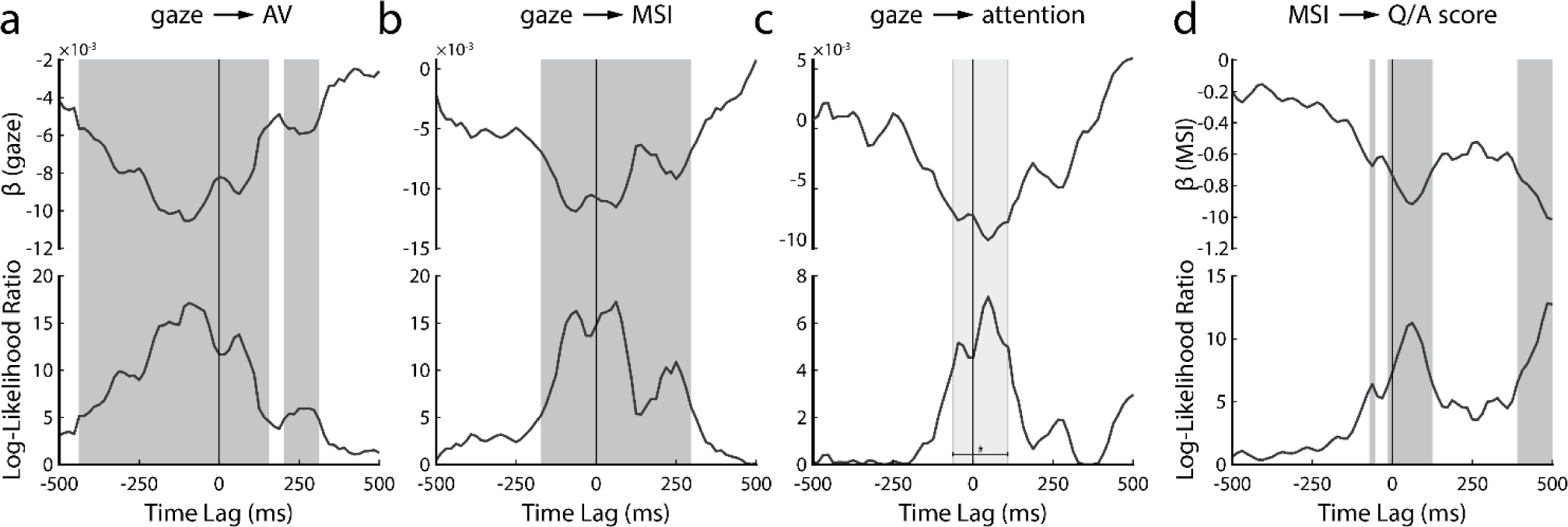
Temporally resolved linear mixed-effect modeling. Upper panel shows the slope parameter of the regression line (β) while the bottom panel shows the Log-Likelihood ratio of the full model over the null model. **a.** Result of predicting attended AV speech tracking from gaze behavior. **b.** Result of predicting multisensory integration of attended AV speech from gaze behavior. **c.** Result of predicting attention benefit on AV speech tracking. **d.** Result of predicting behavioral comprehension scores from corresponding neural measure of multisensory integration. The dark gray bars on a, b and d indicate statistically significant time-lags (*p* < 0.05; FDR-corrected) whereas the light gray bar on c indicates time lags with *p* values less than 0.05, however, the *p* values did not survive FDR correction for multiple comparisons. It is important to note that due to the autocorrelational structure present in both the speech stimuli and the corresponding EEG data, our single-lag reconstruction approach (and by extension, the LME analysis) suffers from some temporal smearing. The relationship between the stimulus and data at a particular lag, say 90 ms, can be quite similar to that at 110 ms, which includes the likelihood of temporal smearing. Data presented in Figures 4 and 5 should therefore be interpreted while keeping this caveat in mind. However, in light of the differences between pre- and post-stimulus decoder weights (Fig. 4a) and the consistency in the multi-modal time course of the gaze effects on different measures (Fig. 5), we feel confident that the analysis is tapping into distinct neural processes and not simply smearing an effect backward in time.

The effect of gaze on multisensory integration is evident for a comparably more restricted time range (Fig. 5b), from −172 ms pre-stimulus to 297 ms post-stimulus. Intuitively, we speculate that this might correspond to the same three neural components discussed above: onset of visual speech, early auditory processing, and an attentional effect. Notably, the very early (−438 ms to −172 ms) effect we saw for speech tracking is absent, suggesting that the impact of gaze in the earliest window is not directly related to the processing of visual speech.

The effect of gaze on attentional benefit (characterized by subtracting the attended and the unattended reconstruction accuracies of AV decoders during the AV trials) is evident for an even narrower time range between −62 ms to 110 ms (Fig. 5c). This range overlaps with the onset of visual speech and the early auditory processing component, as we mentioned earlier in Fig. 5a and b. However, it is important to note that these time-lags did not survive FDR correction (indicated by the light gray bar in Fig. 5c). There was another peak at around 250 ms, likely reflecting attentional effect as we mentioned earlier, however, this peak too was not statistically significant. Therefore, although the peaks in Fig. 5c seem to be consistent with our previous results, we need to be careful about their interpretation since they did not show statistical significance.

Lastly, we were interested in whether the neural processing of visual speech cues could be related to an improved ability to comprehend speech. We fitted a linear mixed-effects model to relate the trial-by-trial listening comprehension scores to their corresponding measures of multisensory integration (i.e., AV – (A+V)). We found that a significant relationship exists for three epochs (Fig. 5d). One epoch coincides with the availability of visual information (∼-100 ms), the second with early auditory processing (∼100 ms), and the third occurs very late (390 ms to 500 ms) and might depend on activity related to semantic processing (Broderick et al. 2018).

## Discussion

How audiovisual integration and selective attention interact with each other remains incompletely understood despite each of them being central to everyday communication. Previous studies have predominantly focused on examining these processes independently, disregarding their interplay. In a previous study, we explored this interaction in the context of natural, continuous audiovisual speech and found a dissociation in an EEG-based measure of multisensory integration for attended vs unattended audiovisual speech (Ahmed et al. 2023). Specifically, we showed that EEG responses to audiovisual speech were best modeled as a multisensory process when speech was attended, but were better modeled as two separate unisensory processes when speech was unattended. In the present study, we aimed to replicate and extend that work in several ways. The primary extension was to explore the question of how attentional modulation of multisensory integration might vary as a function of the amount of visual speech information available to the participants. Using two experiments, we were able to examine this interaction in three different scenarios where participants were — 1) maintaining central fixation, and thereby paying covert attention to one of the two competing audiovisual speakers (experiment 1) 2) directly looking at the face of the target audiovisual speaker while ignoring the other (experiment 2, direct looking condition) and 3) looking at the face of audiovisual speaker they were ignoring, and thereby eavesdropping on the target audiovisual speaker (experiment 2, eavesdropping condition). Our key finding was that our EEG-based measure of multisensory integration was significant only when participants were attending to the corresponding audiovisual speaker *and* when they were directly looking at the face of that speaker.

Our results can be considered through a previously proposed framework wherein listeners extract two distinct types of information from visual speech: correlated and complementary (Campbell 2008; Peelle and Sommers 2015). Correlated information is contained in the visual speech signal itself, wherein the facial movements involved in producing speech sounds exhibit a temporal relationship with the corresponding acoustic waveform. Temporal association is known to exist between the facial movements of the speaker and the speech sound waveform. Notably, previous work has demonstrated that the movements of the mouth area exhibit correlation with the acoustic envelope of the speech across a wide range of frequencies (Chandrasekaran et al. 2009). Various facial features like the eyebrows, jaw, and chin movements have been observed to exhibit correlation with the acoustics as well (Jiang et al. 2002). Studies also indicated that even head movements alone can enhance speech perception, potentially by aligning with the speaker’s voice and conveying prosodic information (Munhall et al. 2004). Visual speech also provides complementary information about the speech that assists in disambiguating acoustically similar speech. These additional visual cues are contained in the visible articulatory detail. Since any articulatory movement made by the speaker is compatible with only a few auditory phonemes, the complementary visual information helps resolve phoneme identity (Peelle and Sommers 2015; Campbell 2008). In the current study, both types of visual information were available in the stimuli, although restricting the gaze behavior of the participants set the extent to which they could utilize each kind of visual information. It is well established that human visual resolution drops off rapidly with distance from the center of vision (Yamada 1969; Loschky et al. 2010), although, people cannot help but notice movement in their peripheral vision (Larson and Loschky 2009; Bayle et al. 2009). As such, in experiment 1, while fixating on the central crosshair, participants were still likely able to utilize the low-level temporally correlated dynamic visual information to help them in solving the cocktail party problem. However, the fact that the speaker’s face was in their peripheral vision likely means that the detailed shape of the speaker’s lips, tongue, and mouth movements were not as fully accessible to the participants, likely leading to a substantial reduction of complementary visual information. This may have caused an overall reduction in the amount of multisensory integration occurring for this central fixation condition. This same is likely true for the eavesdropping condition in experiment 2. Indeed, this is reflected in the fact that our AV decoders did not outperform our A+V decoders for either of these conditions – whether the stimuli were attended or not.

Meanwhile, in the direct looking condition, the availability of both low-level dynamic cues as well as detailed articulatory information produced a significant signature of multisensory integration. Importantly – and replicating our previous study (Ahmed et al. 2023) – this signature of multisensory integration – namely the fact that the AV model outperformed the A+V model – was only true when the speech was attended. Interestingly, this was true despite the fact that, unlike the previous study, the present experiments included an added visual component to the distractor speech (in the previous study, the distractor was purely audio). Furthermore, Ahmed et al. (2023) followed a subject-independent (or generic) modelling approach, while the current study followed a subject-specific approach. Initially, we anticipated that employing a subject-specific design would yield higher reconstruction accuracies; however, we discovered that the reconstruction accuracies were quite similar across the two studies (compare Fig. 3A in the present study to Fig. 2A in Ahmed et al. 2023). This outcome may be attributed to two differences between the experiments used in each study. Firstly, as we mentioned earlier, the previous study introduced an audio-only distractor, while in our current study, the distractor included the corresponding visual speech, requiring participants to suppress both the auditory and the visual conflicting information (which may have received some pre-attentive integration), consequently exposing them to more distractions than may have been caused by an audio-only distractor. Secondly, the study conducted by Ahmed et al. (2023) used different speakers, whereas in our study, we employed different instances of the same speaker. Consequently, speaker identity and voice pitch were not informative cues for facilitating attention, rendering the attention tasks more challenging for participants in the current study. We speculate that the difficulty associated with the attention tasks is reflected in the modest reconstruction accuracies observed in the current study. Nonetheless, both studies are consistent with the finding that multisensory integration is enhanced when participants are attending to an audiovisual speaker that they are also fixating on. Yet, there is one outcome from Ahmed et al. (2023) that we were not able to reproduce in the current study. In Ahmed et al. (2023), the (A+V) decoder was significantly underperforming the AV decoder (i.e., AV < (A+V)) in the unattended AV condition whereas in the current study, we found no significant difference between the performances of the decoders in the unattended conditions. We speculate this may be because the A only and the V only models were derived from audiovisual stimuli different than the audiovisual stimuli they were tested on in the previous study. However, the same audiovisual speaker is used in the current study to both derive and test the models, and this may have established more common information across both AV and (A+V) decoders. It is therefore likely that in the absence of attention, the summation of the unisensory models (A+V) was performing equally to the multisensory AV model.

Due to the involvement of distinct participants in the two experiments (although 7 participants were overlapping), along with variations in experimental aspects such as trial numbers (20 trials in experiment 1 vs 14 trials in experiment 2 per condition), direct group-level statistical comparison across the two experiments became complicated. To address this concern, we combined the data from both experiments and subjected them to linear mixed effects (LME) modeling. Our specific focus was to explore whether gaze could serve as a predictor for various measures (e.g., speech tracking, MSI, attention, behavior scores), all while accounting for individual and experimental discrepancies. Remarkably, our findings affirmed the predictability of these measures by gaze. As gaze diverges from the attended AV speaker, neural tracking of the speaker and its associated multisensory benefit suffers. This interpretation aligns with other findings presented in the paper. With the single-lag analysis, we identified three major time points – consistent with early visual and auditory responses (Karthik et al. 2022; Schroeder et al. 2008), and attentional effects (Patel et al. 2022; Power et al. 2012) – where gaze was associated with stronger speech tracking driven by non-linear multisensory effects. Intriguingly, in a much earlier epoch (−438 ms onwards), speech tracking but not multisensory interactions was affected by gaze. Since we did not expect to see significant stimulus reconstruction so early, it is important to replicate the finding and investigate further. However, our preliminary interpretation is that this relationship does not depend on gaze directly, and is not strictly sensory, but likely represents activity related to predictive processes (Dikker and Pylkkänen 2013; Wang, Kuperberg, and Jensen 2018) that depends indirectly on the visual enhancement of speech processing.

There are also other ways in which the current study can be further developed in future work. We utilized envelope tracking as a speech feature to assess the neural measure of multisensory integration. However, it is important to acknowledge that MSI extends beyond the realm of acoustic processing, and there may be intriguing phenomena occurring at higher levels of linguistic analysis. It would be interesting to investigate how attention and multisensory integration interacts at different hierarchical levels of speech processing and how that interaction varies across gaze behavior. It will also be interesting to derive some visual features of the audiovisual stimuli, such as motion, lip movements, visemes etc. and investigate how the encoding of those features vary depending on gaze and attention. In the future, it would be interesting to utilize intracranial recordings in similar to confirm the timings of the gaze effects on various neural and behavioral measures we sought to explore with our single-lag analysis.

## Author Contributions

FA, AN and EL designed the experiment. FA collected the data. FA and AN analyzed the data. FA wrote the manuscript. FA, AN and EL edited the manuscript.

## Data Availability Statement

The raw data used to generate the findings of this study will be made available by the authors upon request, to any qualified researcher.

## Acknowledgements

The authors would like to thank Dr. Aisling O’Sullivan and Dr. Kevin Prinsloo for providing helpful feedback on the study design, and Lauren Szymula for assisting with data collection.

