## Supplementary Material for "The effect of gaze on EEG measures of multisensory integration in a cocktail party scenario"

To test for multisensory effect in EEG, we compare performance of an AV decoder to a decoder fit on unisensory EEG. Before making our main multisensory comparisons, we wanted to first verify that our decoder models were performing as expected. Using permutation testing, we first verified that the decoders could perform significantly better than chance. Supplementary Figure 1a shows the performances of AV, A and V attended decoders and their null distributions (grey shaded regions – tip to tip of their box and whisker plot) where they all perform better than chance (Supplementary Table 1) in reconstructing the corresponding multisensory or unisensory stimulus envelope (not shown in the figure but the unattended decoders perform better than chance too).

We further found that the V-only decoders perform worse than any other decoders (supplementary Fig. 1a; $AV_d$ vs $V_d$, $p = 4.27 \times 10^{-4}$; $A_d$ vs $V_d$, $p = 0.03$; $AV_c$ vs $V_c$, $p = 0.0084$; $A_c$ vs $V_c$, $p = 0.02$; $AV_e$ vs $V_e$, $p = 0.04$; $A_e$ vs $V_e$, $p = 0.02$; Wilcoxon signed-rank tests). This is not surprising, given that individuals are generally poor lipreaders and the V decoders extract visual information based on its correlation with the acoustic speech envelope as they reconstruct a feature of the speech signal that was not physically present. Importantly, the A-only decoder performed worse than the multisensory AV decoder in the direct looking condition (supplementary Fig. 1a; $AV_d$ vs $A_d$, $p = 0.015$) suggesting that adding the congruent visual speech indeed improves speech representations in the brain. However, this alone is not enough to infer a multisensory integration effect. The presence of visual responses during audiovisual speech can improve reconstruction of the speech signal in the absence of integration. This is why – as we detail below – we quantify multisensory integration by comparing the AV decoders with (A+V) decoders. In line with this issue, it was interesting to note that there was no difference between the performance of the A-only decoders and the AV decoders in reconstructing their respective speech envelopes either in the crosshair fixation or the eavesdropping situation (supplementary Fig. 1a; $AV_c$ vs $A_c$, $p = 0.72$; $AV_e$ vs $A_e$, $p = 0.85$). In these gaze scenarios, modeling the brain response to an attended stimulus using a multisensory audiovisual model is just as good as a unisensory auditory model. We attribute this to the fact that, in the crosshair and eavesdropping conditions, EEG responses to correlated features in the visual speech are much weaker as the participants are not looking directly at the video that corresponds to the audio envelope that we are reconstructing.

We also wanted to validate our (A+V) model to ensure that it was capturing unique sources of information from AV EEG. First, we conducted permutation testing to verify that the (A+V) decoders could reconstruct the envelope of the AV speaker significantly better than chance (Supplementary Table 1). Second, we compared the performance of the (A+V) decoder in reconstructing the envelope of the AV speaker to each of the two unisensory decoders. We found that the (A+V) decoder universally performed better than either A or V models, suggesting that it captures information from both auditory and visual modalities during AV speech processing (attended decoders: $(A+V)_d$ vs $A_d$, $p = 0.015$, $(A+V)_d$ vs $V_d$, $p = 1.22\times 10^{-4}$, $(A+V)_c$ vs $A_c$, $p = 0.001$, $(A+V)_c$ vs $V_c$, $p = 4.38\times 10^{-4}$, $(A+V)_e$ vs $A_e$, $p = 0.003$, $(A+V)_e$ vs $V_e$, $p = 0.002$).

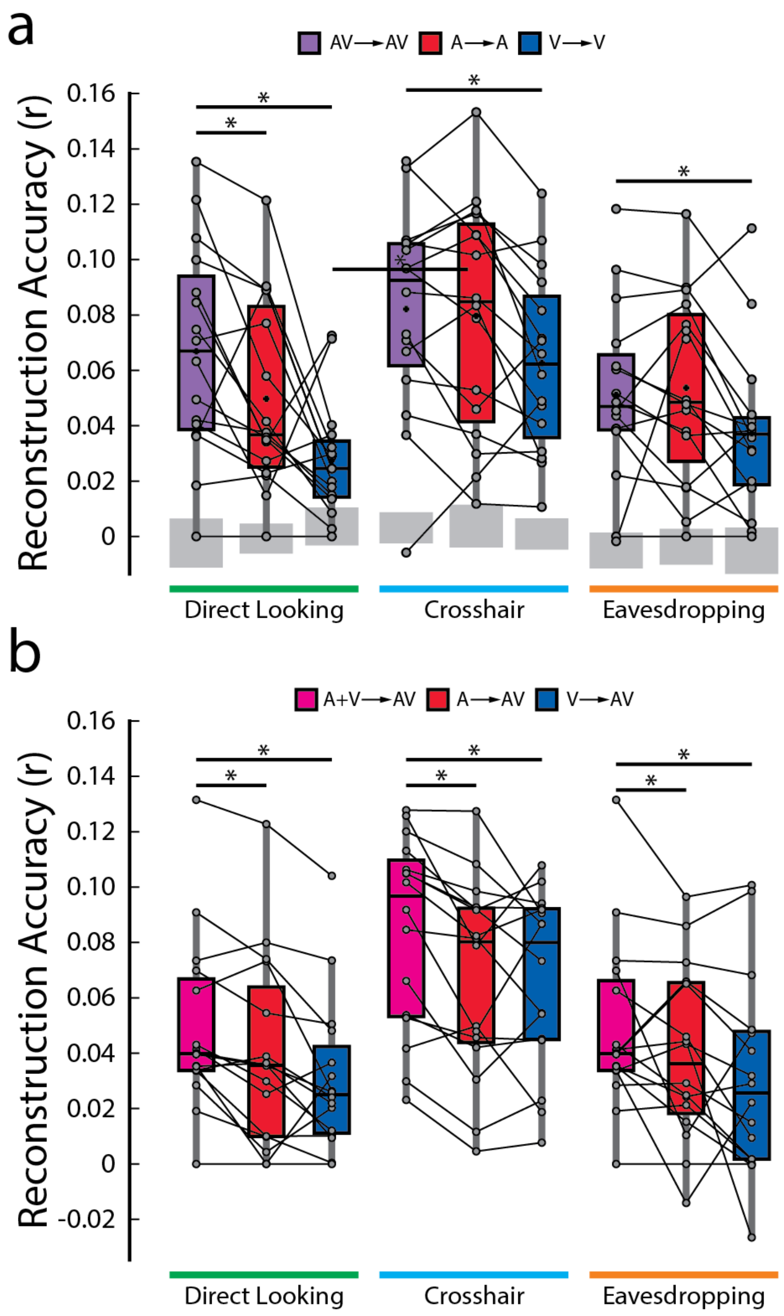


**Supplementary Figure 1.** Decoder performance and validation. **a.** Envelope reconstruction accuracy of each decoder fit to EEG recorded during attention to AV, A, and V speech. Decoder performance is measured on its ability to reconstruct the envelope of left-out trials from its own condition (legend in the form of decoder → condition tested on, e.g., AV→AV, etc.), as is typical for model validation. AV decoders perform better than the unisensory decoders indicating that more speech information is imbedded in AV EEG. **b.** Reconstruction accuracy of the additive-model decoder (A+V) and its constituent decoder models (A, V decoders) as they attempt to reconstruct envelopes from AV speech (A+V→AV, etc.). (A+V) decoder perform better than the unisesnsory decoders, indicating that this model can extract (some of) the multiple sources of information in AV EEG.

**Supplementary Table 1.** Grand-average envelope reconstruction accuracy (r) ± standard deviation. Wilcoxon signed-rank tests against chance level (established by permutation testing) indicate that all decoders could reconstruct the speech envelopes of both attended and unattended audiovisual speakers significantly better than chance.

| Decoders | Attended | Unattended | |
|---|---|---|---|
| $AV_c$ | r = 0.076 ± 0.048; $p = 2.92 \times 10^{-4}$ | r = 0.020 ± 0.035; $p = 2.41 \times 10^{-4}$ | Experiment 1 (N = 16) |
| $(A+V)_c$ | r = 0.079 ± 0.038; $p = 2.41 \times 10^{-4}$ | r = 0.036 ± 0.028; $p = 2.41 \times 10^{-4}$ | |
| $AV_d$ | r = 0.067 ± 0.038; $p = 3.05 \times 10^{-5}$ | r = 0.027 ± 0.021; $p = 2.14 \times 10^{-4}$ | Experiment 2 (N = 15) |
| $(A+V)_d$ | r = 0.049 ± 0.031; $p = 3.05 \times 10^{-5}$ | r = 0.025 ± 0.02; $p = 9.15 \times 10^{-5}$ | |
| $AV_e$ | r = 0.051 ± 0.032; $p = 6.10 \times 10^{-5}$ | r = 0.021 ± 0.017; $p = 9.15 \times 10^{-5}$ | |
| $(A+V)_e$ | r = 0.049 ± 0.033; $p = 3.05 \times 10^{-5}$ | r = 0.021 ± 0.017; $p = 9.15 \times 10^{-5}$ | |
